# Analysis of Single Nucleotide Polymorphisms in human Voltage-Gated Ion Channels

**DOI:** 10.1101/476572

**Authors:** Katerina C. Nastou, Michail A. Batskinis, Zoi I. Litou, Stavros J. Hamodrakas, Vassiliki A. Iconomidou

## Abstract

Voltage-Gated Ion Channels (VGICs) are one of the largest groups of transmembrane proteins. Due to their major role in the generation and propagation of electrical signals, VGICs are considered important from a medical viewpoint and their dysfunction is often associated with a group of diseases known as “Channelopathies”. We identified disease associated mutations and polymorphisms in these proteins through mapping missense Single Nucleotide Polymorphisms (SNPs) from the UniProt and ClinVar databases on their amino acid sequence, taking into consideration their special topological and functional characteristics. Statistical analysis revealed that disease associated SNPs are mostly found in the Voltage Sensor Domain – and especially at its fourth transmembrane segment (S4) – and in the Pore Loop. Both these regions are extremely important for the activation and ion conductivity of VGICs. Moreover, amongst the most frequently observed mutations are those of arginine to glutamine, to histidine or to cysteine, which can probably be attributed to the extremely important role of arginine residues in the regulation of membrane potential in these proteins. We suggest that topological information in combination with genetic variation data can contribute towards a better evaluation of the effect of currently unclassified mutations in VGICs. It is hoped that potential associations with certain disease phenotypes will be revealed in the future, with the use of similar approaches.

## 1. Introduction

Ion channels are transmembrane proteins that enable ion flow via lipid membranes [1]. These proteins govern the electrical properties of membranes of excitable cells and thus are crucial for many physiological functions, such as neuronal signaling, muscle contraction and hormone secretion [2]. Ion channels are mainly classified into Voltage-Gated Ion Channels (VGICs) and Ligand-Gated Ion Channels (LGICs) depending on their activation mechanism [3, 4]. VGICs form a superfamily characterized by the ability to respond rapidly to alterations in membrane potential (as their nomenclature “voltage-gated” suggests), resulting in the selective conductance of ions [5]. The most prominent, voltage-sensitive members of this superfamily can be categorized physiologically into 3 families based on the types of ions that they transport; Voltage-Gated Calcium Channels (Ca_V_s), Voltage-Gated Sodium Channels (Na_V_s) and Voltage-Gated Potassium Channels (K_V_s) [6].

VGICs that belong to these families share some common characteristics. Specifically, they have α-subunits consisting of four domains (DI-DIV). Na_V_s and Ca_V_s consist of one polypeptide chain with four domains [7–9], while K_V_s consist of four polypeptide chains, which assemble to form a functional tetrameric channel [10], which explains the extreme diversity of K_V_s in comparison to the other two families [11, 12]. Each one of these domains shares a common “topological profile” with six transmembrane segments (S1-S6), four of which form the Voltage Sensor Domain (VSD) and a Pore Loop (PL) [7–9]. The PL is located between transmembrane segments S5 and S6 and includes the selectivity filter of the channel [13–16]. The VSD is a modular four transmembrane helix bundle (S1-S4) [17], whose conformational changes upon alterations in membrane potential are responsible for the opening of the channel pore [18–20]. Apart from these distinguished members of the superfamily of VGICs, other families have been described more recently. However, members of these families either do not follow the “topological profile” described above [21–25] or are non-selective, voltage-insensitive and their activation is affected by neurotransmitters and biochemical mediators [24, 26-29].

Each family of VGICs serves a different functional role in the cell. K_V_s mostly control neuronal excitability [30] and participate in solute transport [31], while Na_V_s contribute to the generation of neuronal action potential [32, 33]. Ca_V_s, on the other hand, are involved in a wide range of physiological processes ranging from muscle contraction and neurotransmitter release [34] to gene expression [35]. Due to their versatility, defects in their expression, regulation or function has severe consequences that lead to the emergence of a plethora of diseases [36]. This explains the fact that ion channels currently represent the second largest target group of proteins for drugs in the market, coming immediately after G-protein-coupled receptors [37, 38]. Moreover, advances on compound screening techniques have also designated ion channels as promising targets for the future development of drugs [39].

Progress in the field of human molecular genetics allowed the association between changes in genes coding for VGICs and a group of heterogeneous diseases known as ‘Channelopathies’ [40]. Channelopathies include a broad range of clinical conditions – from epilepsy [41] to cardiac dysfunctions [42] – which arise from gain-of-function or loss-of-function mutations in ion channels [43]. The majority of these disorders are genetically diverse. Same phenotypes arise from different mutations and the same mutation can present with different effects in individuals with distinctive genetic backgrounds [44]. Genetic testing, based on common genetic markers for some of the most common Channelopathies, is commercially available [45]. Some of the most important markers identified for these diseases include Single Nucleotide Polymorphisms (SNPs) [46]. SNPs are inherited single nucleotide substitutions among individuals of a species and are defined as such, when they occur at a rate greater than 1% in a population [47]. Normal substitutions are responsible for genetic variation among the human population. On the contrary, when SNPs result in mutations on regulatory or protein-coding regions of a gene, they can lead to adverse effects and even diseases, such as Channelopathies [46].

The aim of this study was to examine the distribution of missense SNPs and the range of amino acid substitutions in relation to the topology of human VGICs. Our goal was to elucidate the extent to which missense SNPs can be associated with predisposition towards the emergence of a disease. Moreover, we wanted to examine whether the special topological features of VGICs, in combination with genetic data, can aid towards the better characterization of the pathogenic potential of SNPs that are currently unclassified.

## 2. Materials and Methods

### 2.1. Datasets

#### 2.1.1. VGICs dataset

The dataset used in this analysis contains all VGICs that are activated by voltage (voltage-dependent). Their α-subunit forms a functional tetramer, where each monomer comprises of six transmembrane segments and a pore forming loop, responsible for ion transport. From all families of VGICs included in IUPHAR [48], only three meet the aforementioned criteria and are analyzed in this work, namely Voltage-gated Potassium Channels (K_V_s) [49], Voltage-gated Calcium Channels (Ca_V_s) [50, 51] and Voltage-gated Sodium Channels (Na_V_s) [32, 52]. Specifically, all channels used in this study, contain four domains with six transmembrane segments (S1-S6) – where four of them (S1-S4) form the Voltage Sensor Domain (VSD) – and a Pore-loop (PL). Other families characterized as VGICs in IUPHAR are either voltage-independent channels or have a totally different topology than the one described above (see Introduction). The dataset of Ion Channels from IUPHAR and the VGICs entries used in this analysis are shown in Table S1.

#### 2.1.2. Missense SNPs dataset

Human genetic variation data (“Polymorphisms” and “Pathogenic” mutations) were gathered from UniProt database [53] (release date: 10-10-2018) and from ClinVar database [54] (release date: 22-10-2018). A manual mapping between RefSeq Nucleotide Codes from ClinVar to UniProt ACs was conducted (Table S2), in order to gather data from ClinVar only for the isoform that represents the canonical sequence of the VGIC in UniProt. This step was necessary, since position information in UniProt applies only to the “canonical” isoform and incorrect mapping between these two databases leads to numerous mistakes.

The final dataset used in this study emerges from the union of data collected from UniProt and ClinVar. For a more accurate analysis, a common reference code for clinical characterization was used, considering the different terminologies these databases use. Thus, missense SNPs characterized as ‘Disease’ (UniProt), ‘Pathogenic’ or ‘Likely Pathogenic’ (ClinVar) will be referred as “Pathogenic” in the final dataset, while those referred to as ‘Polymorphism’ (UniProt), ‘Benign’, ‘Likely benign’ or ‘Risk factor’ (ClinVar) will be referred simply as “Polymorphism”. Finally, SNPs characterized as ‘Unclassified’ (UniProt), ‘not provided’, ‘not reported for simple variant’, ‘Conflicting interpretations of pathogenicity’, ‘no interpretations for the single variant’ or ‘Uncertain significance’ (ClinVar) will be added to the “Unclassified” dataset. Duplicate entries were merged in one entry in the final dataset, since there were cases where the two datasets overlapped. In cases with conflicting clinical significance for the same entry in the two databases, the clinical significance from ClinVar was adopted (Table S3).

### 2.2. VGICs’ missense SNPs

#### 2.2.1. Mapping of SNPs on biologically significant regions and categorization based on biophysical properties

The determination of three dimensional structures for VGICs, and especially human VGICs, has proven a rather difficult task for the scientific community in the past two decades [55]. The case is similar for the majority of transmembrane proteins [56, 57] and several computational tools have been developed through the years for the prediction of transmembrane segments [58], to provide researchers with a view of the topology of these proteins. Protein databases, like UniProt, have incorporated these data in their membrane protein entries. In this work, data regarding the transmembrane, extracellular and intracellular regions for all VGICs were extracted from UniProt and were transformed, in order to follow the known “topological profile” of VGICs. Specifically, regions were grouped as follows: N-terminal, C-terminal, Intracellular Loops (ILs), Extracellular Loops (ELs), Pore-Loop (PL), Voltage Sensor Domain (VSD) and S5-S6 Transmembrane Regions (S5-S6 TM).

Using this “topological profile”, all “Polymorphisms”, “Pathogenic” and “Unclassified” SNPs were mapped on the sequences of VGICs. Both the exact amino acid substitutions and substitutions grouped in terms of biophysical properties were counted, for the conduction of additional statistical analyses. Asparagine (N), Glutamine (Q), Serine (S), Threonine (T), Tyrosine (Y), Cysteine (C), Methionine (M), Tryptophan (W) and Histidine (H) were considered polar residues, Alanine (A), Glycine (G), Isoleucine (I), Leucine (L), Phenylalanine (F), Proline (P) and Valine (V) were designated as non-polar and Arginine (R), Aspartate (D), Glutamate (E) and Lysine (K) as charged residues [59].

#### 2.2.2. Statistical analysis

Due to the complexity of our data, a simple statistical analysis (e.g. t-test) was not sufficient to examine the parameters we wanted. For this reason, more complex statistical methods were used. Data were subjected to Analysis of Variance (ANOVA [60]) and logistic regression in order to gain an estimate of the correlation between pathological SNPs and their appearance within certain topological regions. More specifically, ANOVA was utilized to get an initial estimate of the variance of appearance of “Polymorphisms” and “Pathogenic” SNPs on transmembrane and non-transmembrane segments. However, ANOVA could not provide sufficient results when the special topological features of VGICs were taken into consideration, and thus, due to the increase of dimensions in our feature set, logistic regression [61] was applied. Regression analysis can aid towards the understanding of how the typical value of a dependent variable changes, when one of the independent variables is adjusted and others are held fixed. During the logistic regression analysis, our focus was on four different topological regions of VGICs. The VSD and PL were examined as the most biologically significant regions of VGICs and the extracellular and intracellular loops due to their role in the interactions between different VGIC domains. Moreover, logistic regression analysis was performed to test the association between the biophysical attributes of the amino acid changes and the pathogenicity status (“Polymorphism” or “Pathogenic”) of missense SNPs.

In order to estimate the statistical significance of all amino acid changes of “Polymorphisms” and “Pathogenic” SNPs in different topological sites, data were subjected to random sampling with replacement (Bootstrap [62]). Bootstrap is a statistical method that allows us to obtain a theoretical estimate regarding the distribution and the standard deviation of data for a given experimentally obtained dataset (amino acid changes in our case). The application of this method on biological data allows an increase in available data in a random and unbiased manner [63]. This analysis was performed on four different starting populations, specifically the samples of a) all pathological SNPs, b) all polymorphisms, c) all pathological SNPs on VSD and d) all pathological SNPs on PL. The method was applied 1000 times for each sample and the average and standard deviation values were determined for all substitutions. The significance of our results was assessed with the calculation of Confidence Intervals (CI) for each SNP in all samples.

Figure 1 shows a workflow of the entire process of data collection and analysis, presented in the previous sections.

**Figure 1.**
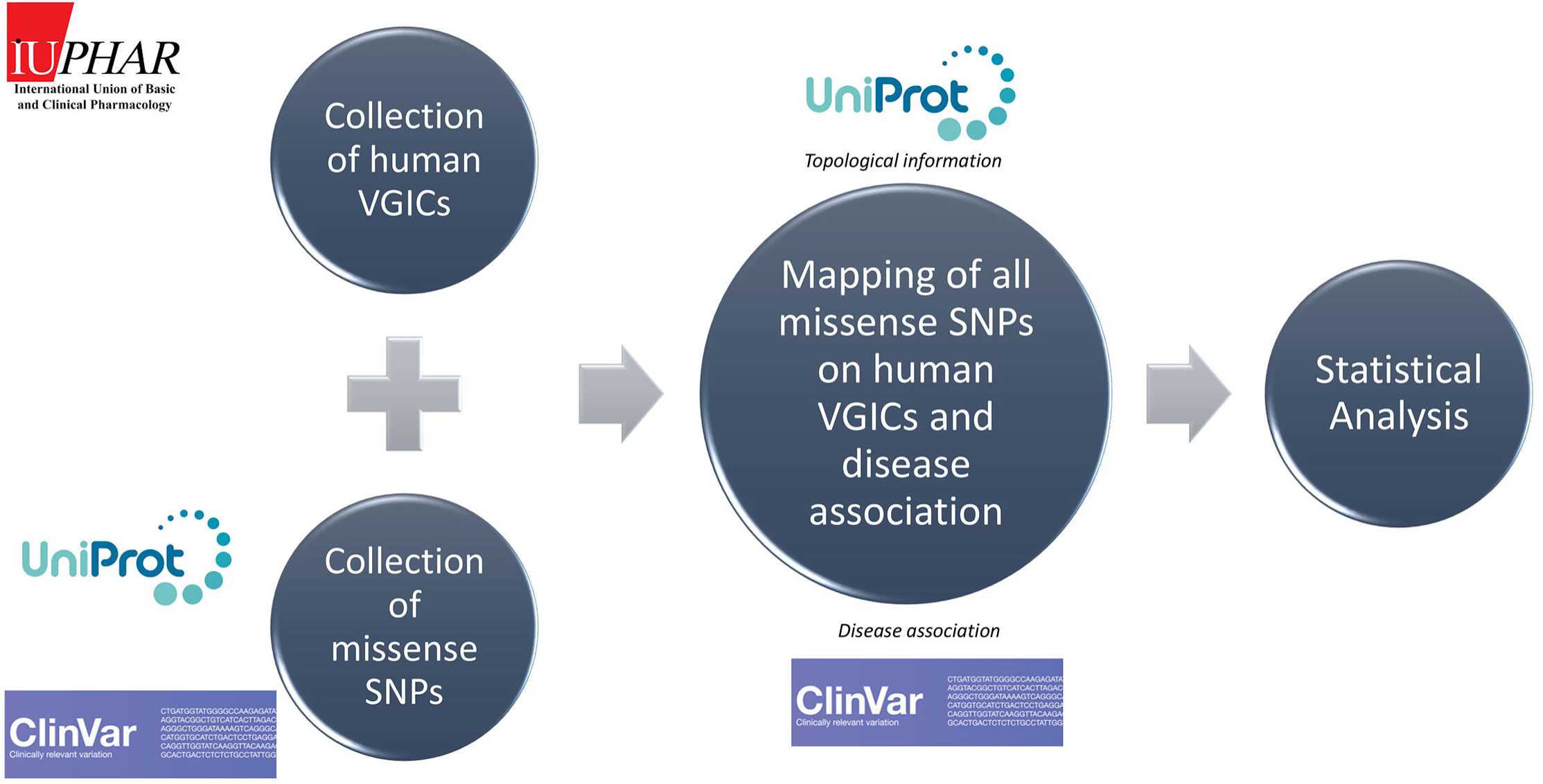
The workflow of data collection and analysis. A dataset of human VGICs was collected from IUPHAR and data regarding missense SNPs of VGICs were gathered from UniProt and ClinVar. These data were manipulated with scripts written in Perl, and all SNPs were mapped on human VGICs in relation to the topological information collected from UniProt. When available, data was isolated from ClinVar regarding the association of specific SNPs with Channelopathies. Statistical analysis was carried out to draw conclusions regarding the significance of our results.

## 3. RESULTS AND DISCUSSION

### 3.1. VGICs’ missense SNPs rates

The dataset of 5311 unique missense SNPs located on 51 out of the 59 known human VGICs was gathered from both UniProt and ClinVar. From those SNPs, 2035 are located on K_V_s, 741 on Ca_V_s and 2535 on Na_V_s (Table S3). Concerning their status, 1765 “Pathogenic SNPs” are found on 33 VGICs, 457 “Polymorphisms” on 42 VGICs and the majority of SNPs are “Unclassified” and account for 3089 SNPs on 41 VGICs (Table S4). The rate of “Polymorphisms” per residue is 5.5×10^-3^, while the rate of “Pathogenic” SNPs per residue is 3.8×10^-2^. This suggests an enrichment of “Pathogenic” SNPs in VGICs and a reduced tolerance to mutations for this protein group, since the appearance of a random SNP in a VGIC usually leads to disease. All SNPs were mapped on VGICs’ sequences and the total numbers per topological region are shown in Table 1. Results in more detail are presented in Table S5. The raw counts of SNPs indicate the VSD, Pore Loop, Intracellular Loops and the C-terminal as the main regions where SNPs appear.

**Table 1.** Total count of SNPs per topological region. The number of SNPs found in each topological region of VGICs.

| *Topological region* | *Pathogenic SNPs* | *Polymorphisms* | *Unclassified* | *Total* |
| --- | --- | --- | --- | --- |
| *N-terminal* | 101 | 48 | 384 | 533 |
| *VSD* | 378 | 31 | 412 | 821 |
| *S5* | 139 | 5 | 104 | 248 |
| *Pore Loop* | 373 | 49 | 397 | 819 |
| *S6* | 178 | 12 | 131 | 321 |
| *Intracellular Loops* | 305 | 142 | 677 | 1124 |
| *Extracellular Loops* | 49 | 24 | 125 | 198 |
| *C-terminal* | 242 | 146 | 859 | 1247 |
| *Total* | **1765** | **457** | **3089** | **5311** |

However, since these regions differ vastly in their length, all data had to be subjected to normalization, based on the length of each topological region. The relative frequencies for each mutation were calculated and are shown in Figure 2. At this point, “Unclassified” SNPs were excluded from any future analysis, since a pathogenicity status could not be assigned for these SNPs. Highest frequencies for both “Polymorphisms” and “Pathogenic” mutations are found in the VSD and ILs, both regions of great biological significance for VGICs. These results were further statistically examined.

**Figure 2.**
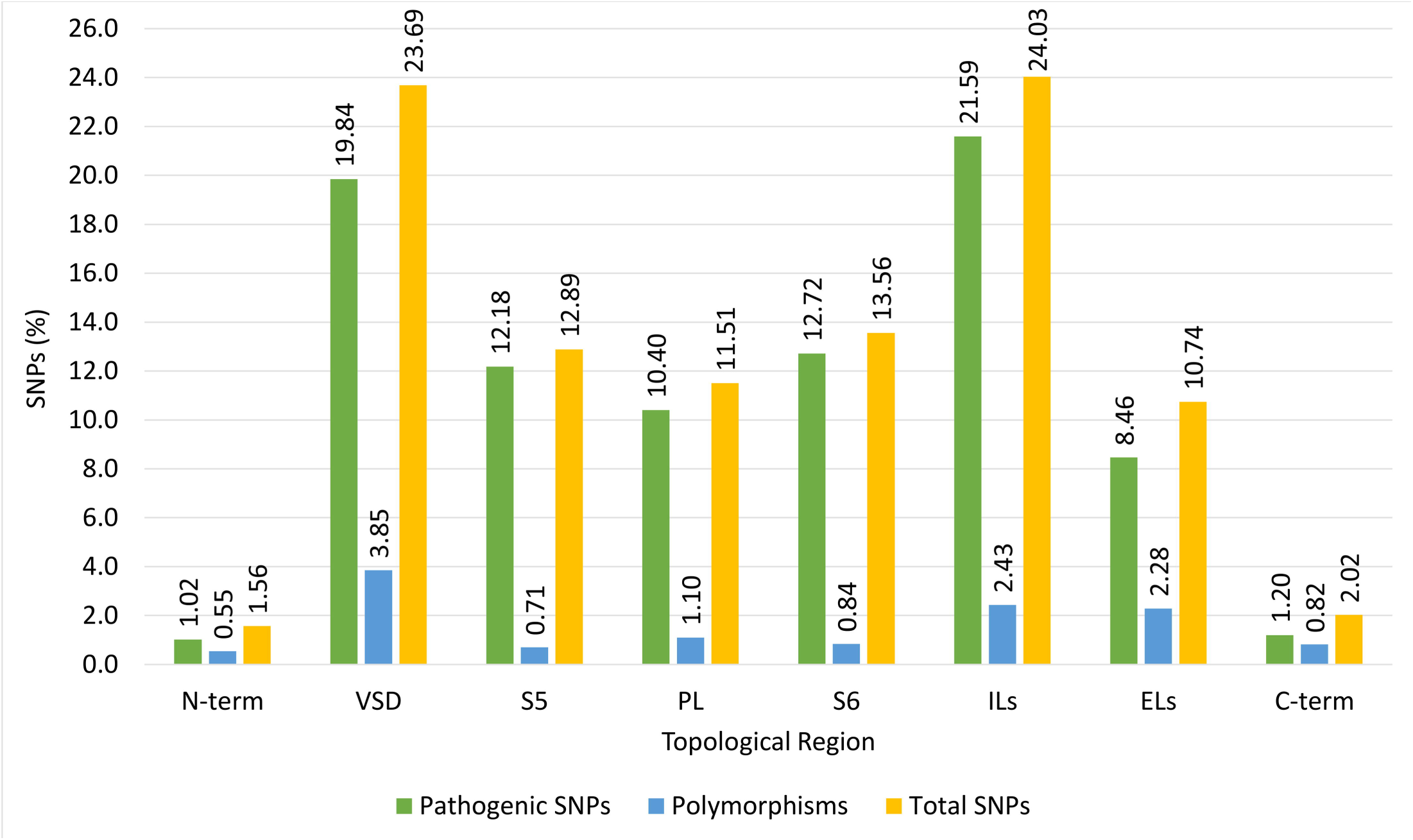
Normalized frequencies of SNPs per topological domain. Green bars represent the frequencies of “pathogenic” SNPs, blue bars show the frequencies of “polymorphisms” and yellow bars the total frequency of “polymorphisms” and “pathogenic” mutations. The VSD and ILs are designated as the most prominent regions for the appearance of SNPs.

### 3.2 Comparison of the distribution of pathogenic mutations’ frequency within topological regions of biological significance

#### 3.2.1. Examining the relation between pathogenicity and transmembrane regions

At first, ANOVA analysis was performed in order to examine the connection between an SNP’s pathogenicity status and its appearance in transmembrane regions. Our null hypothesis was that these parameters are not associated. Results from this analysis showed that the means of observations for “Polymorphisms” and “Pathogenic” mutations differ significantly (Table 2, *P-value<0.05*). However, no difference between transmembrane and non-transmembrane regions was observed and also pathogenicity and transmembrane regions were not found to be related (Table 2).

**Table 2.** Results from two-factor with replication ANOVA. The difference in the distribution of “Polymorphisms” and “Pathogenic” mutations in transmembrane and non-transmembrane regions is examined. The **Sample** difference suggests that the means of observations between “Polymorphisms” and “Pathogenic” mutations varies (*F>F crit, P-value<0.05*). The ***Columns*** difference examines the difference of means between transmembrane and non-transmembrane regions. In this case the null hypothesis cannot be rejected (*F<F crit, P-value>>0.05*), meaning that no difference between SNPs in these two regions could be observed. Finally, the ***Interaction*** difference of means examines the interaction between these two factors (pathogenicity status and appearance in transmembrane or non-transmembrane region). The null hypothesis that there is no interaction between these two factors could neither be rejected (*F<F crit, P-value>>0.05*).. Statistically significant results are shown in **bold**. *SS: Sum of Squares due to the source, df: degrees of freedom in the source, MS: Mean of Squares due to the source, F: F-test statistic, P-value: P-value associated with the F-test, F crit: F critical (from the F distribution)*.

| Source of Variation | SS | df | MS | F | P-value | F crit |
| --- | --- | --- | --- | --- | --- | --- |
| **Sample** | 79853.04 | 1 | 79853.04 | **13.53** | **0.0003** | 3.88 |
| *Columns* | 1.06 | 1 | 1.06 | 0.00018 | 0.98 | 3.88 |
| *Interaction* | 210.4 | 1 | 210.4 | 0.035 | 0.85 | 3.88 |
| *Within* | 1368690 | 232 | 5899.5 |  |  |  |
| *Total* | 1448755 | 235 |  |  |  |  |

#### 3.2.2. Examining the relation between pathogenicity and regions of biological significance

Even though ANOVA can provide a first estimate for the distribution of “Polymorphisms” and “Pathogenic” SNPs on VGICs (Table S6), the method is not sufficient to draw detailed conclusions, regarding the connection between biologically significant regions and pathogenicity. For this reason, logistic regression analysis was carried out, to examine the relationship between the “pathogenicity status” of SNPs and their appearance in specific topological regions. At this point, it should be noted that normalization is not a part of the procedure, since the effect of each independent variable has to be assessed and a normalization process would manipulate the data in such a way that would create bias to the analysis.

Results from the logistic regression analysis of the relationship of pathogenicity status and topological regions are shown in Table 3. *Extracellular and Intracellular Loops* appear as not statistically significant based on our model. As for the extremely statistically significant variables, the *VSD* has the lowest *P-value*, followed by *Other* regions and the *Pore Loop*. This suggests a strong association of these topological regions with the probability of an SNP to be linked with a specific pathogenicity status. The positive coefficient of the estimate in the *VSD* and the *Pore Loop* suggests that all other variables being equal, an SNP in these regions is more likely to be pathogenic. On the contrary, the negative coefficient for *Other* regions indicates that an SNP in these regions is probably not “Pathogenic”.

**Table 3.** Results from logistic regression for pathogenicity status in association with topological regions of biological significance. The distribution of “Polymorphisms” and “Pathogenic” mutations in different topological regions of biological significance is examined. Three regions are statistically significant based on our model. SNPs that are present in the *VSD* and *PL* are more likely to be “Pathogenic” (positive estimate coefficient), while SNPs in *Other* regions of VGICs are probably “Polymorphisms”. Statistically significant results are shown in **bold**.

| *Condition-Region* | *Estimate* | *P-value* | *Significance level* |
| --- | --- | --- | --- |
| VSD | **0.60** | **0.018** | **95%-99%** |
| PL | **0.30** | **0.047** | **95%-99%** |
| Extracellular Loops (ELs) | -0.55 | 0.082 | 90%-95% |
| Intracellular Loops (ILs) | -0.12 | 0.082 | 90%-95% |
| Other | **-0.16** | **0.032** | **95%-99%** |

These results indicate that the VSD and PL are important for VGICs’ functionality. Specific networks of residues that interact with ions in these proteins are extremely important for ion flux regulation and thus, disturbances in these regions often lead to disease [64–69]. More specifically, it has been shown that mutations in the VSD region and especially in the positively charged S4 helix, can cause the pores of these channels to be permanently permeant to ions, at either depolarized or hyperpolarized voltages, where they are normally closed [70].

#### 3.2.3. Examining the interaction between pathogenicity status and the biophysical attributes of mutations

In a different approach, SNPs were grouped based on the biophysical attributes of each amino acid change. The relative frequencies of the entire SNP dataset, as well as, those of “Pathogenic” mutations only, were counted (Table 4). These frequencies demonstrate that the majority of mutations, for all SNPs and for those that are disease associated specifically, are caused from non-polar to non-polar, non-polar to polar and charged to polar changes.

**Table 4.** Relative frequencies of Total and “Pathogenic” mutations in VGICs, grouped based on the biophysical attribute of the residue change. The majority of mutations are from non-polar to non-polar residues, followed by non-polar to polar and charged to polar. The residue changes with the highest frequency for each column are shown in **bold**, those with the second highest with <u>underline</u>, while those with the third highest are shown in *italics*.

| % residue changes | *Total SNPs (%)* |  |  |  | *Pathogenic SNPs(%)* |  |  |  |
| --- | --- | --- | --- | --- | --- | --- | --- | --- |
|  | *VGICs* | *Na<sub>v</sub>s* | *Ca<sub>v</sub>s* | *K<sub>v</sub>s* | *VGICs* | *Na<sub>v</sub>s* | *Ca<sub>v</sub>s* | *K<sub>v</sub>s* |
| polar to polar | 11.25 | 11.34 | 10.10 | 11.55 | 11.22 | 11.63 | 8.47 | 11.08 |
| polar to non-polar | 10.17 | 11.25 | 7.32 | 9.56 | 10.82 | 11.43 | 6.78 | 10.61 |
| polar to charged | 7.16 | 9.05 | 4.88 | 5.05 | 7.71 | 9.54 | 6.85 | 5.20 |
| **non-polar to polar** | <u>16.07</u> | <u>14.64</u> | **20.91** | <u>16.47</u> | *14.94* | <u>14.90</u> | 16.42 | *15.12* |
| **non-polar to non-polar** | **21.29** | **22.25** | *16.72* | **21.51** | **21.50** | **22.01** | <u>17.46</u> | **22.01** |
| non-polar to charged | 8.78 | 7.87 | 9.41 | 9.96 | 9.92 | 8.68 | *16.75* | 10.87 |
| **charged to polar** | *14.81* | *13.20* | <u>19.86</u> | *15.41* | <u>15.03</u> | *13.57* | **20.47** | <u>16.71</u> |
| charged to non-polar | 5.45 | 5.33 | 4.18 | 6.11 | 6.01 | 5.90 | 3.59 | 6.78 |
| charged to charged | 5.04 | 5.08 | 6.62 | 4.38 | 4.86 | 5.00 | 6.35 | 4.51 |

In addition, logistic regression analysis was conducted, to model the relationship of the pathogenicity status and biophysical attributes of the mutations. Results from this analysis are presented in Table 5. In this model, only one case exhibited statistical significance: changes from non-polar to charged residues. The estimate has a positive coefficient (Table 5), suggesting that these changes have a higher probability to lead to the appearance of “Pathogenic” SNPs on VGICs. Despite being on the top three cases with the highest frequencies in our sample, non-polar to non-polar, non-polar to polar and charged to polar mutations were not designated as statistically significant based on our model. This means that even though these mutations appear often, they do not present a higher potential to lead to pathogenicity.

**Table 5.** Results from logistic regression for pathogenicity status in association with biophysical attributes of amino acid changes. The relationship between the pathogenicity status and the biophysical attributes of SNPs is analyzed. Changes from non-polar to charged residues are statistically significant and have a higher probability to lead to disease (positive estimate coefficient). Statistically significant results are shown in **bold**.

| *Condition* | *Estimate* | *p-value* | *Significance level* |
| --- | --- | --- | --- |
| polar to polar | -0.47 | 0.16 | <90% |
| polar to non-polar | 0.10 | 0.71 | <90% |
| polar to charged | 0.34 | 0.41 | <90% |
| non-polar to polar | -0.15 | 0.39 | <90% |
| non-polar to non-polar | 0.00 | 0.99 | <90% |
| **non-polar to charged** | **0.86** | **0.01** | **95%-99%** |
| charged to polar | 0.15 | 0.43 | <90% |
| charged to non-polar | -0.07 | 0.83 | <90% |
| charged to charged | -0.59 | 0.16 | <90% |

#### 3.2.4. Analyzing the distribution of amino acid substitutions for “Polymorphisms” and “Pathogenic” mutations in VGICs

Apart from the analysis of relative frequencies of SNPs based on their biophysical nature, we conducted a more detailed analysis, for each amino acid change. Due to the large size of the tables (20×20) we constructed contour charts to visualize the frequencies of all changes in a concise manner (Figure 3 and Table S7). The contour chart in Figure 3 shows that “Pathogenic” mutations with the higher frequency are those from Glycine (G) to Arginine (R), Leucine (L) to Proline (P), Arginine (R) to Cysteine (C), Arginine (R) to Histidine (H), Arginine (R) to Glutamine (Q), Alanine (A) to Valine (V) and Glutamate (E) to Lysine (K) (Table S7B). From those, E to K, R to H and R to Q mutations are more frequent in the VSD (Table S7C), while R to C and A to V appear mainly in the Pore Loop (Table S7D). All aforementioned mutations can be easily explained based on the standard genetic code, since changes in at least two codons for each substitution are able to induce all these changes.

**Figure 3.**
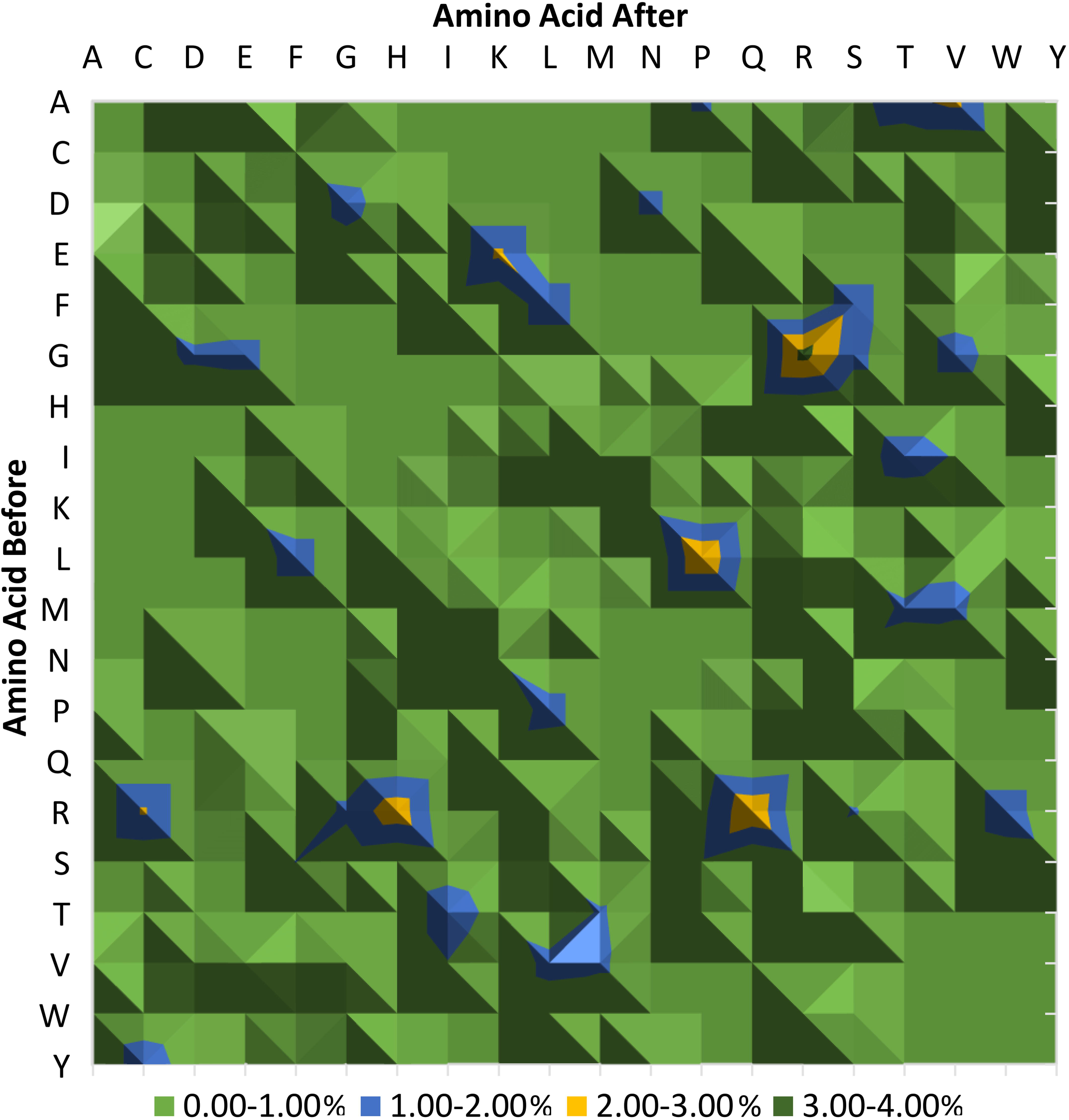
Relative percentage frequencies of “Pathogenic” mutations in the entire length of VGICs. “Pathogenic” mutations with the higher frequency are those in Yellow and Dark Green Peaks in the Contour Chart and specifically G to R, L to P, R to C, R to H, R to Q, A to V and E to K.

In the case of “Polymorphisms”, three mutations emerge as the most frequent, A to T, P to L and R to H (Table S7A).

In an effort to examine the statistical significance of the aforementioned changes, we conducted a random sampling with replacement “Bootstrap” analysis. All observed “Polymorphisms” and “Pathogenic” mutations were counted and were compared to a dataset of mutations produced through 1000 random samplings. This allowed us to increase the available SNP data in order to assess the statistical significance of each change. As shown in Table S8, the most prominent “Pathogenic” mutations, that were identified previously, are all statistically significant.

### 3.3. Focusing on statistically important mutations

Based on our analysis the most prominent disease associated mutations are those of G to R, L to P, A to V, E to K and R to C, H or Q. From those R mutations located on the VSD appear extremely significant.

Previous studies have shown that G to R and L to P mutations in human transmembrane proteins, lead to increased disease susceptibility [71]. VGICs are not an exception, since all such mutations, recorded in transmembrane regions, result to pathogenicity (Table S9A). Most ‘Pathogenic’ G to R mutations are located on the Pore Loops of VGICs. Almost half of them are found on K_V_s (Table S9D). This can perhaps be attributed to the fact that K^+^ selectivity filters contain a signature sequence (TVGYG) [72] with two glycine residues within each loop, and thus a disruption in this sequence can lead to adverse effects for the protein’s functionality. Moreover, when present in transmembrane regions, G to R mutations provide an additional charge in these segments, which leads to the disturbance of the hydrophobic core of these proteins. Even though L and P are both non-polar residues, these mutations are also harmful when located in transmembrane α-helices, since prolines’ side chains provide extra charge due to the exposure of hydrogen bridge acceptors.

In a similar manner, mutations from E to K alter the charge of the mutated residue from negative to positive. This leads to the disruption of the interactions with neighboring residues, explaining the link of such mutations to pathogenicity. Moreover, selectivity filters in the PLs of both Na_V_s (DEKA) [5] and Ca_V_s (EEEE/EEDD) [14, 73] contain E residues and thus such mutations can alter the activation or inactivation mechanism of these channels and the subsequent ion flow. Additionally, Na_V_s [74] alanine mutations and K_V_s [75] A to V mutations in residues adjacent to pore loops have been shown to alter the properties of their selectivity filters, often leading to atypical gating, suggesting an important role of these non-charged residues, as well.

Mutations in VSD have repeatedly been recorded to cause many Channelopathies. A detailed catalogue and analysis of diseases caused by SNPs on VGICs is presented in Section 3.4. Biophysical studies have been carried out to reveal the mechanism underlying these clinical conditions [64, 65, 76, 77]. It has been shown that mutations in the VSD, and specifically in the S4 segment, lead to the emergence of different pathological permeation pathways, known as gating pores [64, 65, 69]. Gating pores have been recorded when the coherence of the hydrophobic septum of VGICs is corrupted in general. These structures activate rapidly, but do not have a mechanism dedicated to their inactivation [70]. This disturbance in ion flow has been associated with many pure electrical disorders, such as Brugada and long QT syndrome [78]. More recently their association with cardiac disorders has also been reported [79]. Mutations in R residues appear to be the key behind the mechanism underlying these disorders [76, 80]. Specifically, S4 arginine neutralizing mutations, such as R to C [81–83], R to H [64] and R to Q [84, 85], have shown different impacts on gating pore currents. Since R residues control the movement of S4 segments, which in turn serve as the cornerstone of voltage sensing, this effect explains the connection of these mutations with diseases such as periodic paralyses [85, 86]. Consequently, gating pores caused by mutations in the VSD could constitute the pathological mechanism of many understudied Channelopathies. Interestingly, these 3 mutations have been designated as the most important arginine mutations in our dataset as well. When present on the VSD they always lead to pathogenicity, stressing the importance of almost another fifty (50) R to C, H or Q SNPs of uncertain significance in the same topological region (Table S9A). The data regarding these mutations is growing and in combination with topological information, potential pathogenic effects can be associated with known or newly identified unclassified SNPs.

### 3.4. Association with Channelopathies

SNPs associated with Channelopathies are found on 33 out of the 59 human VGICs. Some SNPs have more than one association with different diseases. In total 1964 associations were recorded for 1490 unique SNPs and a total of 132 different diseases were identified, excluding general disease categories (Table S10). These diseases can be classified based on the affected human organ system [43]. The majority affects the Nervous System and is mostly associated with Na_V_s (Table 6). Over 65% of SNPs are associated with two major categories of diseases; various types of epilepsies (~35%) and long QT syndromes (~30%) (Table S10A). Furthermore, SNPs of VGICs belonging to different families present a phenotypic overlap, since they lead to the same disorder. For these reasons we decided to look further into each VGIC family and its properties.

**Table 6.** Total count of references to Channelopathies. References from different SNPs are grouped based on the type of disease they are associated with and the VGICs in which they are found. A total of 1490 ‘Pathogenic’ SNPs have 1964 references for 132 Channelopathies. Four major organ systems are affected in Channelopathies; the neuronal, the muscular, the hormonal and the heart muscle system. In the category ‘Other’ SNPs that reference general disease terms (e.g. CACNA-related conditions) and not specific diseases are included. The majority of SNPs is located on Na_V_s and is associated either with Neuronal or Heart Muscle Diseases. ND: Neuronal Disease, MD: Muscular Disease, HMD: Heart Muscle Disease, HD: Hormonal Disease.

| *VGICs (Channels with disease associations)* | *ND* | *MD* | *HMD* | *HD* | *Other* | *Total* |
| --- | --- | --- | --- | --- | --- | --- |
| *Ca<sub>VS</sub> (9/10)* | 101 | 6 | 11 | 5 | 3 | **126** |
| *K<sub>VS</sub> (15/40)* | 286 | 6 | 525 | 0 | 15 | **832** |
| *Na<sub>VS</sub> (9/9)* | 667 | 54 | 270 | 0 | 15 | **1006** |
| ***Total*** | **1054** | **66** | **806** | **5** | **33** | **1964** |

Na_V_s physiological function is the initiation and propagation of action potentials in nerves and muscle fibers [87]. The importance of their inactivation, due to specific mutations, in the development of Channelopathies and particularly neurological disorders, has been thoroughly studied and well established [88, 89], which explains the plethora of data recorded for this type of SNPs in our work. However, it should also be mentioned that this result could be attributed to publication bias, since disorders associated with Na_V_s are those studied mostly by the scientific community [90]. On the other hand, K_V_s are implicated in the establishment of membrane potential and the repolarization after an action potential [87]. A large number of SNPs on K_V_s associated with Heart Muscle Diseases has been recorded via our analysis, in accordance with the effects of the disturbance of their physiological function. Finally, Ca_V_s are responsible for the generation of action potential in the heart and smooth muscles, muscle contraction, neurotransmitter release and intracellular signaling [91]. The biological processes in which they participate, explain why Hormonal Diseases were linked only with a few SNPs on Ca_V_s (Table 6). The small number of representatives for this family though, does not undermine their importance, but rather stresses the need for their future extensive study.

It is obvious that the importance of Channelopathies in the scope of neurological and muscular disorders is gathering growing attention. Recognition of their clinical significance will aid in the profiling of these diseases and the elucidation of their heterogeneity. Moreover, the investigation of rare single-gene pathologies will hopefully provide the appropriate foundation for the delineation of the mechanisms governing more common polygenic disorders, like migraines [88].

## 4. Conclusions

The cascade of data from genome sequencing [92] and Genome Wide Association Studies (GWAS) [93, 94] has enabled the identification of novel associations between SNPs and specific disease phenotypes [95]. However, concerns have been raised in the past about many SNPs discovered through GWAS. They may have auxiliary roles in the development of a disease phenotype and thus turn out to be unsuitable candidates for the improvement of health care through genetic testing [96, 97]. For this reason, efforts are being made so that current analytical protocols can shift into more integrated approaches that account for the genomic context of genotype-phenotype relationships [98]. Bioinformatics could take a more active role in the address of this issue.

In this work, we focused on VGICs, a protein family with great pharmacological significance. We have proved that ‘structural’ protein information, like the topological profile of VGICs can contribute to the evaluation of the implications of genetic variations in this protein group. Our results showed that mutations that alter the charge and structural properties of amino acid chains are extremely important in the development of pathogenicity. Furthermore, two regions have been designated as the most important topological regions for disease implication; the VSD and the PL. More specifically, arginine (R) mutations in the S4 helix of the VSD appear to be the most significant disease associated mutations, based on our analysis and in accordance with literature data. We hope that the integration of topological information – when available – in the biostatistics protocols, used to catalog SNPs associated with specific disease phenotypes, will aid in the identification of clinically relevant mutations and in the determination of the pathogenic status of the numerous available unclassified SNPs.

## Supporting information

## Acknowledgements

The authors thank the National and Kapodistrian University of Athens for support.

---

**Table S1: The dataset of Voltage Gated Ion Channels from IUPHAR**. Green Color in Column AC indicates the Voltage dependent ion channels incorporated in the final dataset (namely KVs, CaVs and NaVs). The column “Comment” explains briefly the reason why a channel or an entire family of channels (orange cells) is excluded from our analysis (for more details please refer to the main manuscript).

**Table S2: Mapping between RefSeq Nucleotide Codes and UniProt ACs** for the canonical isoforms of VGICs. This mapping was used to gather data from ClinVar regarding the SNPs of VGICs in the GRCh38 version of the proteome.

**Table S3 (A): The full dataset of missense SNPs** located on KVs (2035 SNPs), CaVs (741 SNPs) and NaVs (2535 SNPs). The excel sheet named “Full” contains a dataset of unique SNPs from both databases (5311 SNPs). **(B): The dataset of missense SNPs** located on KVs (669 SNPs), CaVs (188 SNPs) and NaVs (1128 SNPs) as isolated from UniProt. The excel sheet “UniProt” contains all SNPs isolated from UniProtKB (1985 SNPs). **(C): The dataset of missense SNPs** located on KVs (1964 SNPs), CaVs (593SNPs) and NaVs (1732 SNPs) as isolated from ClinVar. This excel sheet named “ClinVar” contains all SNPs isolated from ClinVar (4289 SNPs).

**Table S4**: **The dataset of Voltage Gated Ion Channels with SNPs**. In the sheet “Pathogenic SNPs” 33 VGICs with 1765 SNPs are shown. In the sheet “Polymorphisms” 42 VGICs with 457 SNPs are shown and in the sheet “Unclassified” 41 VGICs with 3089 SNPs are shown.

**Table S5**: **The full dataset of missense SNPs located on KVs (2035 SNPs), CaVs (741 SNPs) and NaVs (2535 SNPs).** The topological domain in which each SNP is located is given. Topological domains were characterized based on the topological profile of VGICs and are the following: C-terminal, IL1-11: Intracellular Loop 1-11, EL1-8: Extracellular Loop 1-8, S1-S6: Transmembrane Segment 1-6 (for K_V_s), S(1-6)D(I-IV): Transmembrane Segment 1-6 from Domain I-IV (for Ca_V_s and Na_V_s), PL: Pore loop (for K_V_s), PL1-4: Pore loop 1-4 (for Ca_V_s and Na_V_s), N-terminal.

**Table S6: Results from two-factor with replication ANOVA.** The difference in the distribution of “Polymorphisms” and “Pathogenic” mutations in regions of biological significance (VSD, PL, ILs, ELs, Other) is examined. The Sample difference suggests that the means of observations between “Polymorphisms” and “Pathogenic” mutations varies (F>F crit, P-value<0.05). The Columns difference examines the difference of means between the different regions of biological significance, and suggests that there is no difference observed between the appearance of SNPs in these regions (F<F crit, P-value>0.05). Moreover, the Interaction difference of means between these two factors (pathogenicity status and appearance in biologically significant regions) could not be rejected (F<F crit, P-value>0.05), and suggests that these factors are not associated. Statistically significant results are shown in bold. SS: Sum of Squares due to the source, df: degrees of freedom in the source, MS: Mean of Squares due to the source, F: F-test statistic, P-value: P-value associated with the F-test, F crit: F critical (from the F distribution).

**Table S7**: **Relative frequencies of amino acid substitutions.** The relative frequencies for “Polymorphisms” and “Pathogenic” Mutations for the entire VGICs (Sheets: *Polymorphisms_Total* and *Pathogenic_Total*) are given, along with the relative frequencies for regions with biological significance (*Pathogenic_VSD* and *Pathogenic_PL*).

**Table S8**: **Bootstrap analysis.** The significance of the different kind of substitutions. Tables represent the observed (OBS) values, the average values of random sampling of 1000 times (AVG), the standard deviation of the random sampling (SD), the upper (CIHI) and lower (CILO) limits of the 95% Confidence Intervals (CI) and the statistical significance testing for each observation (TEST). The result is considered statistically significant if the observed value lies within the upper and lower limits of the 95% confidence intervals. Amino acids before substitutions are shown in rows and amino acids after substitutions in columns. Four tests were performed. One for all “Polymorphisms” (*Polymorphisms_Total*), one for all “Pathogenic” mutations (*Pathogenic_Total*), one for “Pathogenic” mutations in the VSD (*Pathogenic_VSD*) and one for “Pathogenic” mutations in the Pore Loop (*Pathogenic_PL*).

**Table S9**: Counts of the 7 most prominent mutations identified in VGICs and Na_V_s, Ca_V_s and K_V_s, grouped based on the topological regions they are located.

**Table S10**: **(A) SNPs with disease associations**. In total 1964 associations were recorded for 1490 unique SNPs. Some SNPs have more than one association with different diseases. ND: Neuronal Disease, MD: Muscular Disease, HMD: Heart Muscle Disease, HD: Hormonal Disease, **(B) List of Channelopathies VGICs' SNPs are associated with.** A total of 132 different diseases were identified (excluding general disease categories), **(C) List of Common Channelopathies between different VGIC families**.

## Abbreviations

VGICs: Voltage-Gated Ion Channels
LGICs: Ligand-Gated Ion Channels
SNPs: Single Nucleotide Polymorphisms
Ca_V_s: Voltage-Gated Calcium Channels
Na_V_s: Voltage-Gated Sodium Channels
K_V_s: Voltage-Gated Potassium Channels
PL: Pore Loop
VSD: Voltage Sensor Domain
ILs: Intracellular Loops
ELs: Extracellular Loops
ANOVA: Analysis of Variance
GWAS: Genome-Wide Association Study

